# A PhyloFisher utility for nucleotide-based phylogenomic matrix construction; *nucl_matrix_constructor.py*

**DOI:** 10.1101/2023.11.30.569490

**Authors:** Robert E. Jones, Erin P. Jones, Alexander K. Tice, Matthew W. Brown

## Abstract

Phylogenies built from multiple genes have become a common component of evolutionary biology studies. Molecular phylogenomic matrices used to build multi-gene phylogenies can be built from either nucleotide or protein matrices. Nucleotide-based analyses are often more appropriate for addressing phylogenetic questions in evolutionarily shallow timescales (i.e., less than 100 million years) while protein-based analyses are often more appropriate for addressing deep phylogenetic questions. PhyloFisher is a phylogenomic software package written in Python3. The manually curated PhyloFisher database contains 240 protein-coding genes from 304 eukaryotic taxa. Here we present *nucl_matrix_constructor.py*, an expansion of the PhyloFisher starting database, and an update to PhyloFisher that maintains DNA sequences. This combination will allow users the ability to easily build nucleotide phylogenomic matrices while retaining the benefits of protein-based pre-processing used to identify contaminants and paralogy.

## INTRODUCTION

Multi-gene phylogenetics, phylogenomics, has revolutionized the way we understand the evolutionary history of life (Delsuc et al., 2005, Duchêne 2021, Lozano-Fernandez 2022). For deep evolutionary relationships, such as the origin of eukaryotes or the diversification of animals, it is common practice to use protein-based phylogenomic analyses (Burki et al., 2020; Simion et al., 2017). This is because protein sequences are much less prone to homoplasy compared to DNA sequences because of the slower rate of observable evolution and the larger alphabet (20 amino acids compared to 4 nucleotides) (Philippe et al., 2011, Kapli et al., 2023). This makes it helpful to use amino acid sequences rather than DNA sequences for ortholog and paralog determination. However, the usefulness of DNA sequence based phylogenetics for resolving relationships between closely related species is well documented, recent work suggests their utility in deep phylogenetics too (Kapli et al., 2023).

PhyloFisher is a Python3 based phylogenomic software package. PhyloFisher includes a manually curated database of 240 protein-coding genes from 304 eukaryotic taxa covering known eukaryotic diversity, a novel tool for ortholog selection, and utilities that will perform diverse analyses required by state-of-the-art phylogenomic investigations (Tice et al., 2021). Previously, only the predicted protein sequences were included in the PhyloFisher software package. We have now expanded the PhyloFisher starting database to include the corresponding nucleotide sequences for the 240 protein coding genes present in the database. This expanded database also comes with an update to PhyloFisher. v1.3.1 that will encourage users to supply nucleotide sequences that correspond to the protein sequences from taxa added to PhyloFisher. This allows users to easily create nucleotide-based matrices to use for phylogenomics analyses.

To demonstrate a use case for the tool, we perform phylogenetic analyses of the Saccharomycetaceae clade of budding yeasts using 2 different gene sets as in Tice et al. (2021) but inferred through nucleotide based phylogenomics. The first gene set is made up of the top 10% of trees with the highest Relative Tree Confidence Scores from the BUSCO1292 dataset of Shen et al., (2018), as in Tice et al., (2021). These RTC scores were calculated and binned by the PhyloFisher utility, *rtc_binner.py*. The second dataset is made up of the 240 genes in the starting PhyloFisher database.

## METHODS

### Implementation

*nucl_matrix_constructor.py* is written entirely in Python3. The utility takes the output of *prep_final_dataset.py*, which contains amino acid sequences for each gene, and a tab separated values file (TSV) with paths to coding sequence (CDS) files as input. The script begins by building BLAST (Camacho et al., 2009) databases using MAKEBLASTDB for each CDS file. TBLASTN (Camacho et al., 2009) is then performed with the CDS as the database and the amino acid sequence for each gene for each taxon as the query. The TBLASTN results are then parsed and the nucleotide sequence, which corresponds to the amino acid sequence query, is collected. The collected nucleotide sequences are then placed into files, one for each gene. The nucleotide sequence files are then aligned using MAFFT (Katoh & Standley, 2013) with the -- auto option. The resulting alignments are then trimmed by trimAl (Capella-Gutiérrez et al., 2009) with a gap threshold of 0.80. The trimmed alignments are next concatenated into a supermatrix. Two files, indices.tsv and occupancy.tsv, are produced. The former contains the gene positions in the super matrix, and the latter contains taxa occupancy for each gene.

### Operation

The recommended method to install PhyloFisher is simply via a Conda environment. Please see https://thebrownlab.github.io/phylofisher-pages/getting-started/installation/ for installation instructions. Additionally, if the user prefers, PhyloFisher can be run inside a Docker container. The PhyloFisher Docker container can be obtained at https://quay.io/biocontainers/phylofisher. We recommend running *nucl_matrix_constructor.py* in a high-performance computing (HPC) environment. The utility was successfully run with 9 taxa and 158 genes on Rocky Linux v8.8 with 40 threads in 0.55 CPU hours. Instructions on how to include nucleotide sequences into existing databases can be found on the PhyloFisher website (https://thebrownlab.github.io/phylofisher-pages/).

## USE CASES

To demonstrate the application of *nucl_matrix_constructor.py* we performed phylogenetic analyses of the Saccharomycetaceae clade of budding yeasts. We analyzed two datasets. The first gene set is made up of the top 10% RTC scoring trees of the Shen et al. BUSCO1292 dataset. These RTC scores were calculated and binned by the PhyloFisher utility, *rtc_binner.py*. The second dataset is made up of the 240 genes in the starting PhyloFisher database. We built phylogenomic trees from nucleotides sequences of the two data sets. This was accomplished by first running *nucl_matrix_constructor.py* once the prep_final_dataset step of the PhyloFisher workflow was reached. The DNA sequence phylogenies were built using IQ-TREE2 v2.2.0.3 (Minh et al., 2020) and through RAxML (Stamatakis, 2014) under the model GTR+G+F. Overall the phylogenomic trees are consistent with the results of Shen et al. 2018 and Tice et al. 2021. The ML tree built from the top 10% RTC scoring trees of the Shen et al. 2018 BUSCO1292 dataset display the same topology as the protein dataset of Tice et al. 2021. However, with nucleotides the MLBS support is higher (100%) compared to protein dataset (93%). Interestingly, the PhyloFisher 208 ortholog dataset built with nucleotides shows a different topology than PhyloFisher 208 ortholog dataset built with proteins. The topology of the ML concatenation-based tree built with nucleotides has the TYV clade (*Tetrapisispora, Yueomyces*, and *Vanderwaltozyma*) breaking up the SNKN clade (*Saccharyomyces, Nakaseomyces, Kazachstania*, and *Naumovozyma*), and the SNKN clade forming a paraphyly.

## CONCLUSIONS

Here we present a simple but useful utility to construct nucleotide-based phylogenomic matrices. This resource allows users to perform nucleotide-based phylogenomic analyses utilizing the PhyloFisher (Tice et al., 2021) methodology easily. Up to this point, PhyloFisher only allowed for protein-based analyses. Thus, *nucl_matrix_constructor.py* will allow for the expansion of PhyloFisher into sub-fields of phylogenomics where nucleotide-based analyses are more appropriate than protein-based.

## ACKNOWLEDGEMENTS

This work was supported by the United States National Science Foundation (NSF) Division of Environmental Biology (DEB) grant 2100888 (http://www.nsf.gov) awarded to MWB.

---

**Figure 1:**
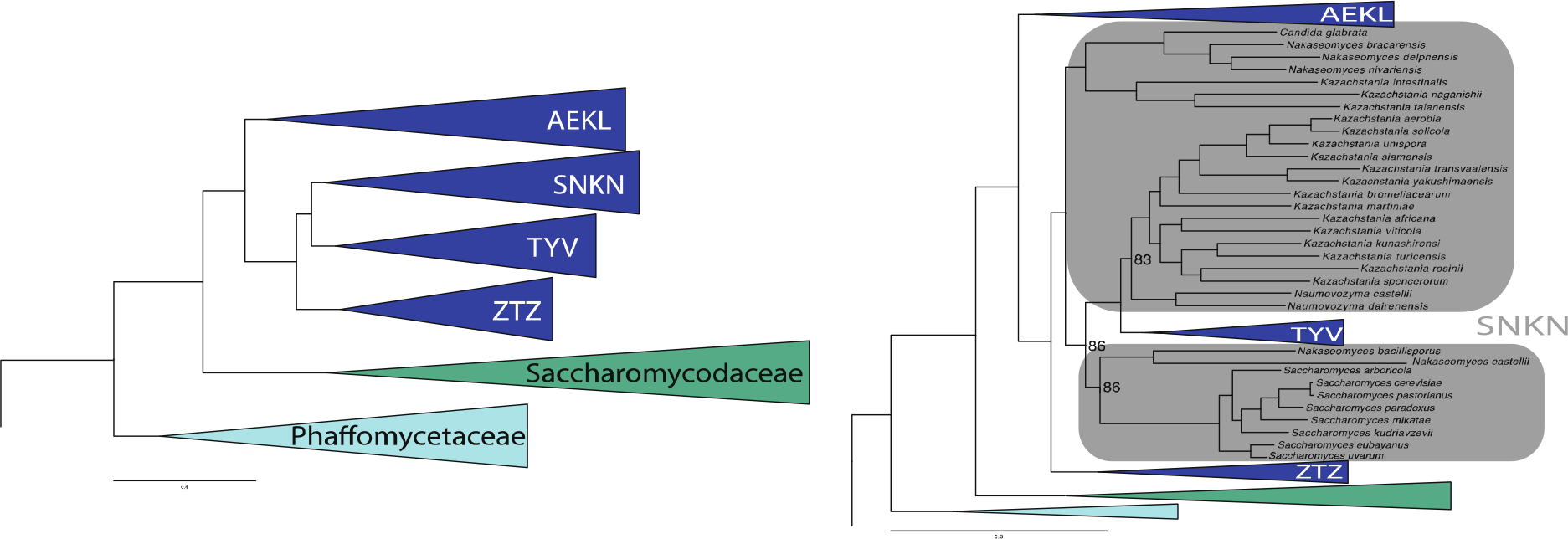
Phylogenetic reconstruction of the tree of Saccharomycetaceae using DNA sequences. Maximum likelihood trees were built using IQ-TREE2 v2.2.0.3 (Minh et al., 2020) under the model GTR+G+F utilizing a DNA sequence matrices from two datasets. The first gene dataset, shown on the left, is made up of the top 10% RTC scoring trees of the Shen et al., (2018) BUSCO1292 dataset as presented in Tice et al. (2021). These RTC scores were calculated and binned by the PhyloFisher utility, *rtc_binner.py*. The second dataset, shown on the right, is made up of the 240 genes in the starting PhyloFisher database. Sub-clades that make up the Saccharomycetaceae are shown in dark blue which are comprised of AEKL (*Ashbya, Eremothecium, Kluyveromyces*, and *Lachancea*), SNKN (*Saccharyomyces, Nakaseomyces, Kazachstania*, and *Naumovozyma*), TYV (*Tetrapisispora, Yueomyces*, and *Vanderwaltozyma*), and ZTZ (*Zygosaccharomyces, Torulaspora*, and *Zygotorulaspora*) clades, while the outgroup clades of the Saccharomycodaceae and the Phaffomycetaceae are shown in dark green and cyan, respectively.

## Notes

https://thebrownlab.github.io/phylofisher-pages/

### Competing Interest Statement

The authors have declared no competing interest.

